# Depriving mice of sleep also deprives of food

**DOI:** 10.1101/2021.12.21.473656

**Authors:** Nina Đukanović, Francesco La Spada, Yann Emmenegger, Guy Niederhäuser, Frédéric Preitner, Paul Franken

**Affiliations:** Center for Integrative Genomics, University of Lausanne, Lausanne, Switzerland; Mouse Metabolic Evaluation Facility, Center for Integrative Genomics, University of Lausanne, Switzerland

**Keywords:** Circadian rhythms, Clock genes, Sleep deprivation, Food intake, Energy expenditure, Body composition, Lean body mass, Fat

## Abstract

Both sleep-wake behavior and circadian rhythms are tightly coupled to energy metabolism and food intake. Altered feeding times in mice are known to entrain clock-gene rhythms in brain and liver and sleep-deprived humans tend to eat more and gain weight. Previous observations in mice showing that sleep deprivation (SD) changes clock-gene expression might thus relate to altered food intake and not to the loss of sleep *per se*. Whether SD affects food intake in the mouse and how this might affect clock-gene expression is, however, unknown. We therefore quantified i) the cortical expression of the clock genes *Per1, Per2, Dbp*, and *Cry1* in mice that had access to food or not during a 6h SD, and ii) food intake during baseline, SD, and recovery sleep. We found that food deprivation did not modify the SD-incurred clock-gene changes in the cortex. Moreover, we discovered that although food intake during SD did not differ from baseline, mice lost weight and increased food intake during subsequent recovery. We conclude that SD is associated with food deprivation and that the resulting energy deficit might contribute to the effects of SD that are commonly interpreted as a response to sleep loss.

## Introduction

One presumed important adaptive advantage of an endogenous, circadian time-keeping system is that it enables the organism to anticipate the daily changes in the environment such as the availability of food. The circadian system thus ensures that the animal is awake and active before food becomes available and, along with other constraints, defines the animal’s temporal ecological niche. In conflict experiments, in which food availability is restricted to a time-of-day when the animal is normally asleep and does not engage in feeding behavior, such as during the light period in the case of a nocturnal rodent, the animal’s behaviour and physiology must adjust to anticipate the new feeding regimen. Studies in mice and rats show that these feeding-related adjustments are not immediate and it can take several days until variables such as wakefulness and corticosterone production become entrained and peak prior to the new time of food availability [1–4]. This entrainment is also evident at the molecular level. In cortex and liver, the phase of the oscillation in the expression of circadian clock genes adjusts to that of the new feeding time, thereby dissociating from clock-gene rhythms in the central circadian pacemaker in the suprachiasmatic nuclei (SCN), which remain entrained to the light-dark cycle [3, 5, 6].

The expression of clock genes is also modified when sleep is altered. For instance, keeping animals awake at a time-of-day when they normally sleep, affects clock-gene expression in tissues peripheral to the SCN including the cerebral cortex, liver, and kidney [7–11]. Because clock-gene expression responds to alterations in both feeding and sleep-wake behavior, it has been suggested that sleep deprivation (SD) may alter clock-gene expression indirectly, through increased food intake. Although plausible, this notion is less straightforward than it seems. First, the effects of SD on clock-gene expression are acute and can be observed already after 3h of SD [8, 10], which is inconsistent with the gradual re-entrainment over several days observed in food-restriction experiments. Second, while humans do eat more (and gain weight) when sleep deprived [12–14], it is unclear whether and, if so, how SD affects food intake acutely in the mouse. The aim of the current study was therefore two-fold. We first sought to establish whether access to food during SD affects the SD-induced changes in clock-gene expression in the cerebral cortex. We then assessed whether, like humans, mice eat more when kept awake. We found that during SD mice ate as much as during matching baseline hours and that the cortical clock-gene expression changes after SD were unaffected by food deprivation. Moreover, despite normal food intake, mice lost weight during the SD and ate more during the subsequent recovery, demonstrating that the SD incurred a metabolic imbalance. We believe this imbalance to result from the metabolic cost of staying awake which we illustrate by putting food-intake into a sleep-wake context.

## Results

### Sleep deprivation affects clock-gene expression independent of food intake

To determine whether the SD-induced changes in the cortical expression of clock genes are caused by changes in food intake, we compared expression changes in the three core clock genes *Per1, Per2*, and *Cry1* and the ‘clock-controlled’ gene *Dbp*, in mice with and without access to food during a 6h SD starting at light onset (ZT0). Expression of the activity-induced short isoform of *Homer1, Homer1a*, considered a reliable marker of sleep homeostatic drive [15, 16], was quantified in parallel. The mRNA levels reached after SD were contrasted to non-SD controls with or without access to food that were sacrificed at the same time-of-day (ZT6).

In mice with access to food, SD elicited a robust increase in *Homer1a, Per1, Per2*, and *Cry1* expression and decrease in *Dbp* expression (**Fig. 1**), similar to what was previously shown [17]. Depriving mice of food during the SD did not modify these responses (**Fig. 1**), demonstrating that increased food intake does not contribute to SD’s effect on clock-gene expression.

**Figure 1:**
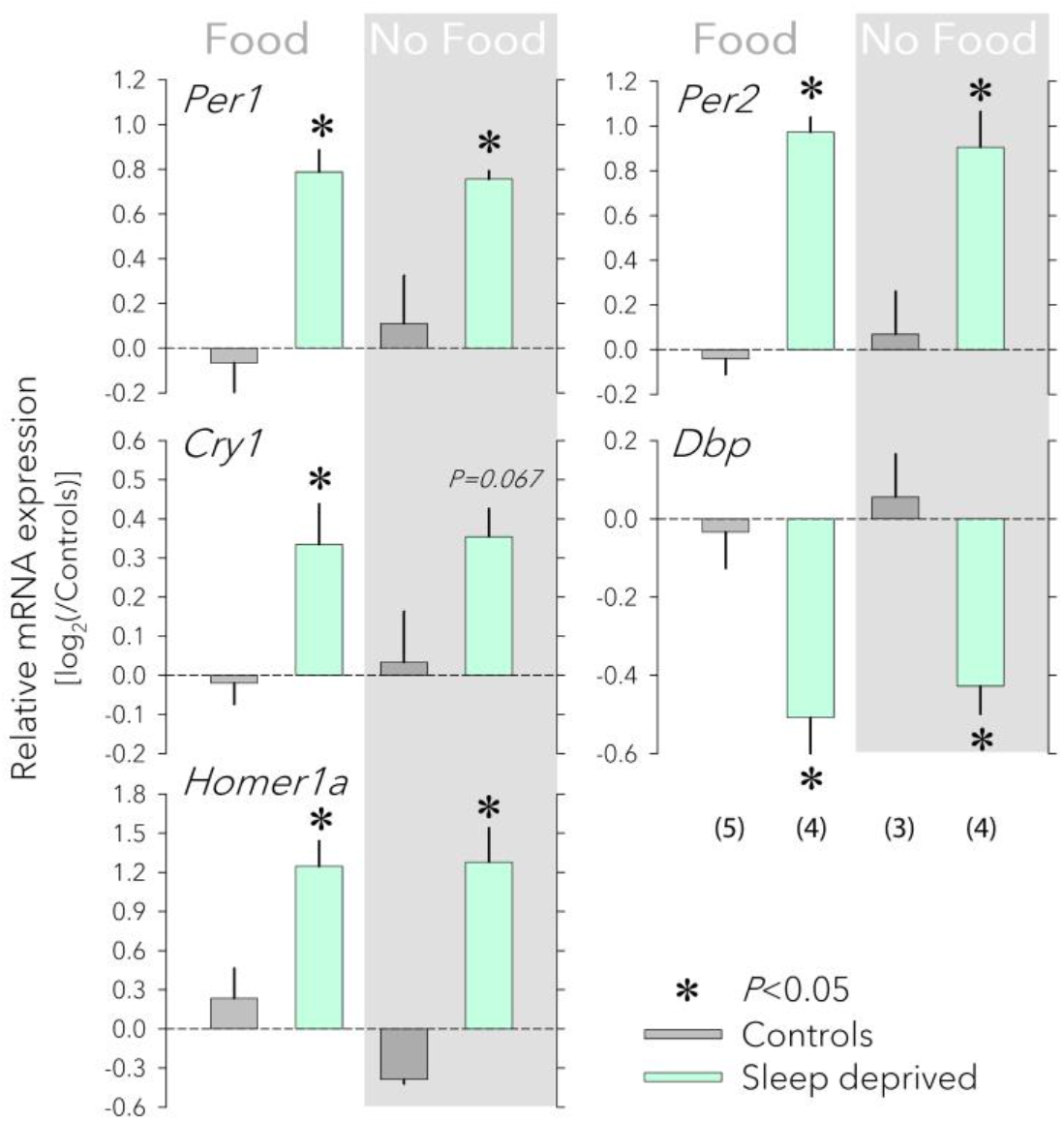
Effects of food availability on cortical clock-gene expression. Having access to food (‘Food’, left) or not (‘No Food’, right, grey background) during a sleep deprivation (SD) did not affect the SD-induced changes in the cortical expression *Per1, Per2, Cry1, Dbp*, and *Homer1a*.Values for each transcript were first expressed relative to the expression of three housekeeping genes (*Eef1a, Gapdh*, and *Rsp9*) within each sample and subsequently to the mean of the two control groups (0 level, dark-grey bars) that each accompanied the SD (mint bars) of the two food conditions. Number of mice for each group in parentheses (error bars reflect 1 SEM). Food did not change the SD response (2-way ANOVA factor ‘Food’ p>0.57, factor ‘SD’ p<0.0021; interaction p>0.17 for each of the 5 transcripts). Asterisks designate a significant SD effect within each food condition (2-sided post-hoc *t*-tests; *p*<0.05). mRNA levels of housekeeping genes were not affected by either SD or food access (not shown). All tissues were collected at ZT6.

### Sleep deprivation does not alter food intake yet animals eat more during recovery

The idea that altered food intake might contribute to the SD-induced changes in clock-gene expression is based on the assumption that animals eat more during the SD. To test this assumption, we measured food intake using the *Phenomaster* feeding monitoring system, under baseline conditions (5 days prior SD; Days −4 to 0), during a 6h SD (Day 1, ZT0-6), and during recovery (Day 1, ZT6-24 and Days 2 to 5; see **Fig. 2**).

**Figure 2:**
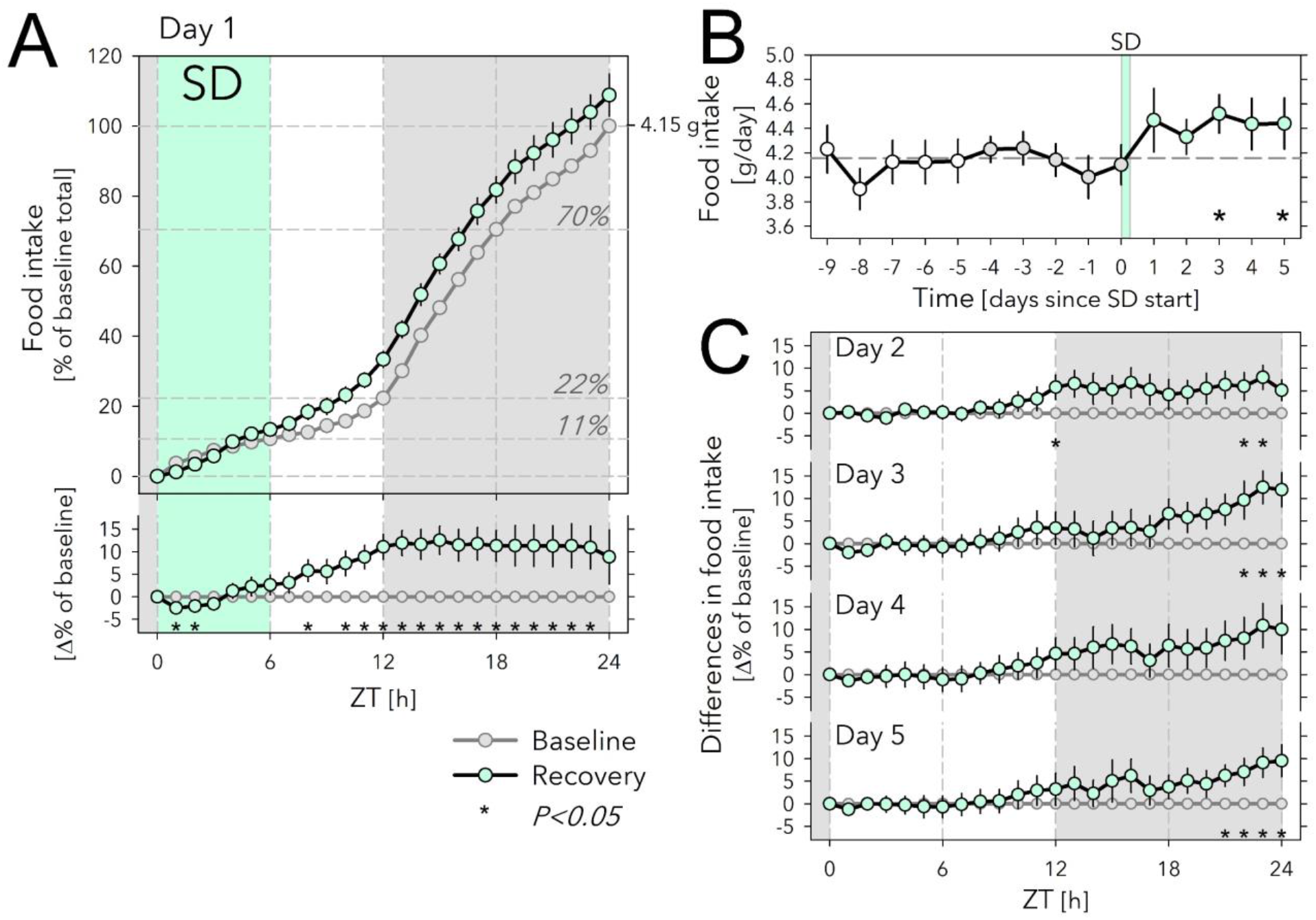
Effects of sleep deprivation on food intake. (**A**) Hourly values of food-intake accumulated over 24h and expressed as % of average daily food intake during the last 5 baseline days (Days −4 to 0; 100% = 4.15g) preceding the sleep-deprivation (SD) day (Day 1). Food intake during SD (mint symbols and area) did not differ from baseline (grey symbols, 5-day averages). Horizontal lines in graph indicate % of total daily food eaten at consecutive 6h intervals during baseline. Lower graph: SD-baseline difference in accumulated food-intake curves depicted in the upper graph. Gray area indicates the 12h dark period. Asterisks designate hours with significant differences in food-intake from baseline (paired 2-sided *t*-tests; *p*<0.05; *n*=11-17; see Methods for *n*/day). (**B**) Daily food intake for the 10 days before and the 5 days after the SD. White symbols (Days −9 to −5) depict values for 5 additional baseline days recorded in cohort 1 (*n*=6). Please note that these extra baseline days were not included in the analyses. Horizontal dashed line reflects mean over Days −4 to 0 (=100% in A). SD affected food-intake (Days 1 to 5 *vs*. Days −4 to 0; 2-way ANOVA factor ‘SD’ *p*=0.004, ‘Day’ *p*=0.89, interaction *p*=0.78). Asterisks mark days with significant increase in food-intake compared to baseline (average Days −4 to 0; paired 2-sided *t*-tests; *p*<0.05; *n*=11-17; see Methods for *n*/day). (**C**) Recovery-baseline differences in accumulated hourly food-intake values for Days 2 to 5. As on Day 1, the increase in food intake on Day 2 was mainly due to extra food eaten during the 2^nd^ half of the light phase, while for Days 3 to 5 extra food intake occurred at the end of the dark phase. Asterisks as in A. Error bars mark ±1 SEM.

Under baseline conditions, mice ate 4.15g chow per day, most of which was consumed during the 12h dark phase (**Fig. 2A**). Nevertheless, mice ate substantial amounts during the light phase as well, reaching 22.3% of daily food intake, half of which was consumed during the first 6h (ZT0-6); i.e., the time the SD was scheduled on Day 1 (**Fig. 2A**).

Surprisingly, while over the first 2h of the SD mice ate somewhat less compared to matching baseline hours, when calculated over the entire 6h SD, the amount consumed did not differ from baseline (13.3 *vs*. 10.7%; **Fig. 2A**). Given this lack of an SD effect, it was perhaps even more surprising that over the first 6h of recovery (Day 1, ZT6-12) they ate 73% more than baseline (20.1 *vs*. 11.6% of baseline daily food intake; **Fig. 2A**) so that over the entire 12h light phase, that included the SD, mice ate 0.45g more than during baseline (+48%; 1.38g *vs*. 0.93g). This increase in accumulated food intake was maintained throughout the 12h dark period of Day 1 because mice ate the same amount of food as in the preceding baseline dark periods. As a result, by the end of Day 1, SD mice had eaten 4.51g; i.e., 9% more than the average daily food intake (**Fig. 2A**).

Remarkably, daily food uptake remained upregulated over the next 4 recovery days (Days 2 to 5), although statistical significance was not reached for Day 4 (**Fig. 2B,C**). Interestingly, the time-of-day the extra food was consumed changed over the 5 recovery days; while during Day 2, similar to Day 1, animals ate more in the second half of the light phase, on subsequent days this occurred towards the end of the dark periods (ZT21-24 on Days 3 to 5; **Fig. 2C**). During the first 9h of the light periods (ZT0-9) food intake was remarkably constant among Days 2 to 5 and did not deviate from baseline (**Fig. 2C**).

### Sleep deprivation dissociates food intake from time-spent-awake

Knowing that extended wakefulness represents a metabolic burden [18–20], combined with our current observation that mice eat more after the SD but not during the SD, suggests that depriving mice of sleep incurred a metabolic deficit or, in other words, that per unit of time-spent-awake during SD, food intake was insufficient. To investigate this further, we quantified the relationship between time-spent-awake and food intake during baseline, SD, and recovery sleep. While for technical reasons we could not record EEG/EMG and food intake simultaneously in the same individuals, we have abundant normative sleep-wake data of mice of the same inbred strain, sex, and age submitted to the same experimental protocol and conditions, to allow group-level comparisons. Here we used published data of Diessler, Jan *et al*. [21] and Hor, Yueng *et al*. [22].

In baseline conditions, the time course of mean hourly values of food intake was tightly correlated with the time-spent-awake accounting for 86% of the variance in food intake (**Fig. 3A,B**; adjusted R^2^=0.86; *p*<0.0001, *n*=24 1h intervals). Moreover, the residuals of the linear regression did not systematically vary with time-of-day (**Fig. 3A**, lower graph), suggesting that mice ate the same amount of food per unit of time awake, independent of time-of-day.

**Figure 3:**
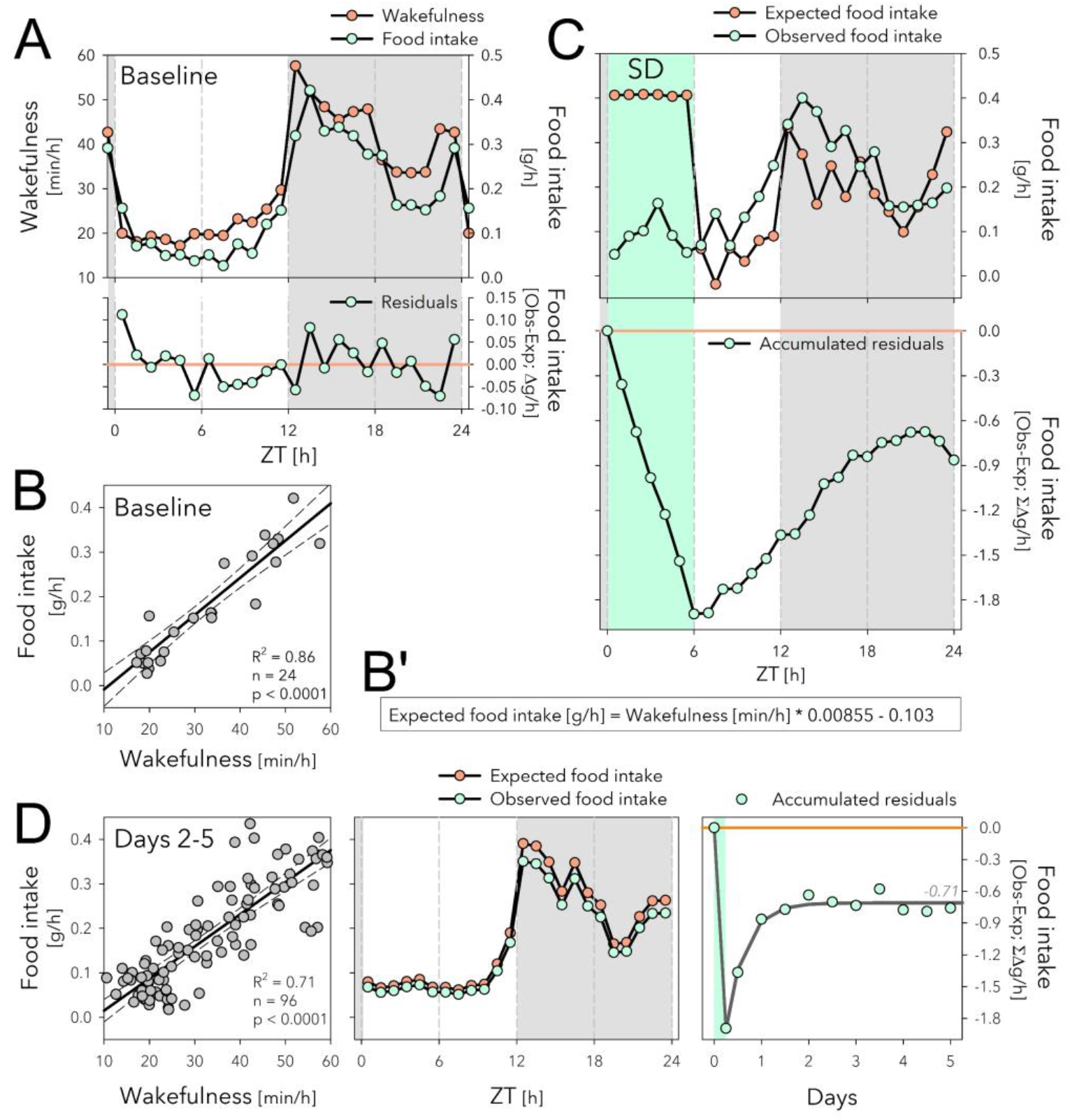
The sleep-deprivation (SD) evoked dissociation between food intake and time-spent-awake might underlie increased food intake during recovery. (**A**) The time course of hourly values of food intake in baseline (mint symbols; mean over Days −4 to 0; same data as in Fig. 2A), closely follows that of time-spent-awake (orange symbols). (**B**) Mean hourly values of wakefulness and food intake strongly correlated, with only small residuals that did not systematically vary with time-of-day (lower graph in A). (**B’**) Equation of the linear regression in panel B used to predict *expected* food intake in panels C and D (middle) based on time-spent-awake. (**C**) Given the equations in B’, the *expected* food intake can be calculated for Day 1 including the SD (upper graph). SD induced a disassociation between food intake and time-spent-awake and animals accrued a deficit of 1.89g of food intake during the SD that reduced to 0.86g by the end of Day 1 (lower graph). (**D**) During Days 2 to 5 after SD, the relationship between waking and eating was highly correlated as in baseline (left panel, compare to B), and wake-derived *expected* food intake (given equation B’) predicted remarkably well the observed data (average over the 4 days, middle panel). The 0.86g food deficit estimated for Day 1 (see C), was maintained during the following 4 days and stabilized at −0.71g (right panel; non-linear regression, exponential saturating function fitted to 12h values).

Using the relationship between food intake and time-spent-asleep established for baseline (**Fig. 3B’**), we then predicted expected food intake based on the sleep-wake values observed for Days 1 to 5 (**Fig. 3C,D**). This analysis revealed a profound dissociation between time-spent-wake and food intake during SD; while mice are expected to eat 0.41g when awake for 60min in baseline conditions (**Fig. 3A,B’**), the observed average hourly food intake during SD was 0.09g; i.e. 4.6-fold lower than expected (**Fig. 3C**, upper graph). The discrepancy between expected and observed food intake amounts to a total deficit of 1.89g food intake when accumulated over the 6h SD (**Fig 3C**, lower graph). This deficit of feeding relative to time-spent-awake was reduced to 0.86g by the end of Day 1 because during the 18h of recovery mice both ate more (+0.52g; **Fig. 2A**), and slept more (+2.04h) than during the corresponding baseline period (**Fig. 3C**, lower graph).

During subsequent recovery days (Days 2 to 5), the baseline relationship between food intake and time-spent-awake was re-established and the wake-derived expected food intake predicted remarkably well the observed values (**Fig. 3D**, left and middle panels). During Days 2 to 5, the 0.86g loss of food intake estimated for Day 1, did not further substantially decrease and stabilized at around 0.71g (**Fig. 3D**, right panel). This despite the increase in food intake accumulated over these 4 days (+1.15g; **Fig. 2B**), perhaps in part because mice slept less (−89min, corresponding to 0.66g extra food required according to the equation in **Fig. 3B’**).

### Sleep deprivation causes weight loss

The above analyses indicate that during SD mice eat less than expected based on the time-spent-awake. This could represent an energy imbalance likely to underlie the increase in food intake we observed during the 6h following the SD. The ‘eating-less-than-expected’ phenomenon could also lead to weight loss during the SD, which would provide more direct evidence documenting an energy deficit. To assess this, we measured each animal’s weight at the end of a 6h SD ending at ZT6 and compared this to its baseline body weight reached at the same time-of-day measured 1 and 4 days prior to the SD. All animals were given *ad libitum* access to food. We found that SD indeed led to a highly-significant 4.8% loss in body weight (−1.3±0.1g; paired 2-sided *t*-test; *p*=1.3E-5; *n*=9; **Fig. 4A**), despite mice having free access to food.

**Figure 4:**
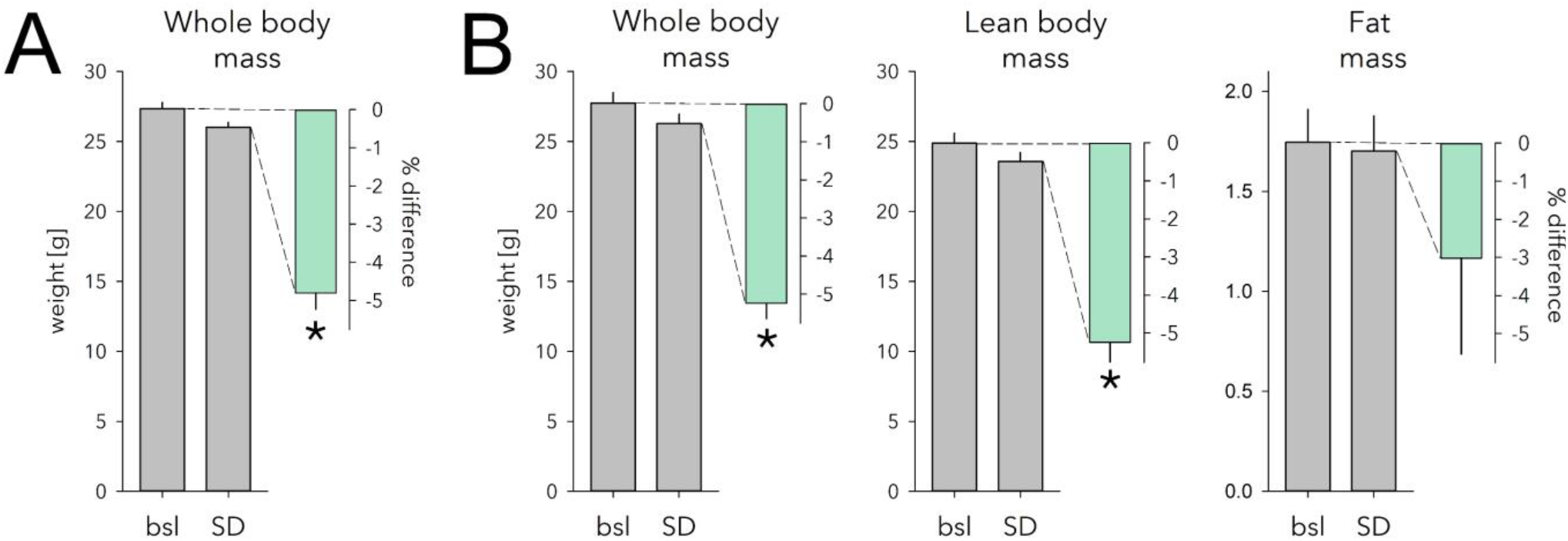
Effect of sleep deprivation (SD) on body weight and body composition. (**A**) SD caused a 4.8% weight loss in mice with *ad libitum* access to food (mint bar; *p*=1.3E-5, paired 2-sided *t*-test, *n*=9). Body weight measured after the 6h SD (grey bar) ending at ZT6 was 1.3±0.1g lower compared to that reached at this time-of-day on two preceding baseline days (bsl; grey bar; average of the 2 days). (**B**) Body composition was measured in a separate cohort following the exact same protocol as in A. SD similarly reduced whole body weight in this cohort (−1.5±0.1g, 5.2%; *p*=9.6E-5, paired 2-sided *t*-test, *n*=6). Lean body mass decreased to the same extent (−1.3±0.1g, 5.2%; *p*=3.1E-4). Fat mass was reduced as well but not significantly so (−0.04±0.04g, 3.0%, *p*=0.33). Details as in A. * marks significant SD *vs*.bsl effects. Note different scale for grams of fat mass.

To identify which body compartment (fat or lean body mass) accounted for this decrease in body weight, we next assessed the effect of SD on body composition in a separate cohort of mice. Consistent with the previous two cohorts, SD mice lost 5.2% body weight compared to baseline (−1.5±0.1g, *p*=9.8E-5, **Fig. 4B**). This decrease was due mainly to a 5.2% loss of lean body mass (−1.3±0.1g, *p*=3.1E-4; **Fig. 4B**). Fat mass was reduced as well, but not significantly so (−0.04±0.04g, 3.0%, *p*=0.33; **Fig. 4B**). Water loss did not seem to play a role as tissue hydration was not affected by SD (baseline: 85.3±0.3% *versus* SD: 84.8±0.2%; *p*=0.23; see Methods).

## Discussion

The first main finding of our study is that having access to food or not during SD does not affect the SD-induced changes in cortical clock-gene levels. This indicates that changes in clock-gene expression during SD do not relate to increased food intake as observed in SD studies in humans. The second main finding is that, contrary to expectation, SD deprives mice of food. While eating as much as during corresponding baseline hours and despite having free access to food, animals nevertheless lose 5% of their body weight when kept awake and eat more than baseline during the first 6h of subsequent recovery. These observations show that the amount of food eaten during SD was insufficient to cover energy expenditure leading to a metabolic deficit.

### Sleep deprivation or food deprivation?

As mice have a small body size they have high metabolic rates compared to humans [23, 24]. C57BL/6J mice, at the body weights seen in our study (27-28g), spent 51-57kJ/day [25]. This energy expenditure (EE) has to be balanced by energy intake through food, which, estimated on the 4.15g/day we observed and given the caloric content of the food (see Methods), amounts to 54kJ/day. Corrigan *et al*. [25] also quantified the daily changes in EE, which as for food intake (**Fig. 3**), matched the distribution of our wakefulness data remarkably well (**Suppl. Fig. 1**). From the relationship between hourly values of time-spent-awake and EE one can estimate that the baseline time-spent-awake (i.e., ca. 13h/day), compared to sleeping 24h, adds 15kJ; i.e., 27% of the daily EE.

Mice lost ca. 1.4g body weight during the 6h SD of which 1.3g was lean body mass and 0.04g fat. Using energy density estimates for fat and lean-body mass (39.5 and 7.6kJ/g, respectively [26]), the 1.4g weight loss we observed would produce 11.5kJ used to offset the cost of being kept awake while not eating enough. Adding the energy from food intake during the SD (7.2kJ), the available energy during the SD would have amounted to 18.7kJ. According to the time-spent-awake to food-intake correlation, mice were expected to eat 1.89g more food during the SD than they did, 1.03g of which was recovered by eating and sleeping more. Using these estimates, the incurred energy deficit would amount to 13.5-24.8kJ, a range compatible with the above estimation based on the tissue energy estimates.

A similar, 1.2g (5%) loss of body weight was observed after a 5h food deprivation (FD) at approximately the same time of day (ZT23-4) [27]. In that study, the weight loss was due to a decrease of 0.2g fat mass and of 0.8g of lean body mass which would represent 14kJ of energy. Although these remain back-of-the-envelope calculations and the actual metabolic cost associated with an SD should be assessed experimentally, both the FD and SD study do demonstrate that short-term decreases in food intake at time-of-day animals normally eat little, can have important consequences on physiology and behavior. Besides weight loss, both interventions share a number of other physiological consequences such as increased levels of corticosterone and amylase and metabolites such as carnitines [21, 28–31]. Sleep was not quantified in the FD study and it is therefore not possible to assess whether its effects are solely due to reduced energy intake or also because of increased energy expenditure resulting from increased time-spent-awake related to food-seeking behavior and/or stress [32, 33]. Similarly, the consequences of SD that sleep researchers attribute to the incurred sleep loss, might instead be due to deficits in food intake. For example, the time-spent-awake to food-intake correlation illustrates how the well-known rebound in sleep time after SD can be put into an energy context as, together with eating more, it appears an efficient measure to offset the SD-incurred energy deficit.

We found food intake to be increased, compared to baseline, during the days after the SD, pointing to potential long-term effects on energy balance. Alternatively, the slight decrease in sleep-time over these 5 recovery days and/or normal growth might have contributed to higher energy intake. We previously reported on long-term effects on the brain transcriptome notably on the expression of circadian clock genes [22]. Given that clock genes and energy metabolism are tightly coupled together [34, 35], these observations point to a hereto unnoticed and surprisingly long-lasting effects on energy metabolism after only a single, short-term perturbation of sleep. Long-lasting electrophysiological consequences of a 6h SD have also been observed in the rat hypothalamus, including the arcuate nucleus [36], which is important in regulating food intake and energy balance [37].

### Circadian, sleep-wake, or food driven?

An obvious and intriguing question is why mice allow body weight to drop instead of simply eating more during the SD. This is also unexpected because acute SD in the rat signals hunger considering the rapid increase in ghrelin during SD, especially in the hypothalamus [38]. Circadian factors and/or the presence of light might actively inhibit food intake during the rest phase. Food restriction experiments in which access to food is restricted to this inappropriate circadian phase indeed show that mice only slowly adapt to such paradigm and initially lose weight [39]. The increase of food intake observed in the remainder of the light phase (ZT6-12) demonstrates, however, that mice, when hungry, are able to eat more at this circadian phase and in the presence of light. The increased food intake at this time is all the more surprising as it competes which sleep’s homeostatic drive which is considered to be greatest immediately after the SD. During the subsequent dark phase, the circadian phase when mice habitually eat the most, food intake no longer differed from baseline but the rebound in sleep time was largest [40]. This increase in sleep time can also be seen as a behavioural measure to balance the energy budget. Alternatively, as the SD protocol is considered a mild stressor [41], stress might have kept mice from eating more.

The circadian clock, located in the SCN, drives the daily changes in the sleep-wake distribution. Also, food intake and EE are considered to be under strong and direct circadian control [35, 42]. Our correlation analyses demonstrated, however, that the changes in food intake result from changes in time-spent-awake as food-intake expressed per minute of wakefulness does not appreciatively vary with time-of-day. Similarly, EE closely tracks time-spent-wake [43] (**Suppl. Fig. 1** [25]). This tight relationship between waking, EE, and food intake, makes sense given the high metabolic rates of mice because, as we show here, the extra energy spent by being awake requires immediate refuelling to avoid deficits in energy balance.

Clock genes engage in transcriptional-translational feedback loops (TTFL) that constitute the molecular time-keeping circuitry underlying circadian rhythm generation [44]. Besides being part of the circadian TTFL circuitry, several core clock genes act as sensors of environmental and systemic signals [5, 45–49]. For example, the clock gene *Per2* can be rapidly induced by stress, light, changes in temperature, and blood-borne systemic cues [50–54]. As a consequence, in brain tissues peripheral to the SCN such as the cerebral cortex, *Per2* expression is determined by both circadian and sleep-wake driven factors [11]. Given our current results, the acute SD-incurred changes in clock-gene expression clearly do not result from eating more. Instead, it can be argued that clock-gene expression in the cortex responds to the rapid changes in metabolic drive associated with wakefulness both under baseline and SD conditions. In this context, the increase in *Per2* expression observed under conditions of food restriction might be associated with the waking preceding to the time window of food accessibility window (i.e., food anticipatory activity) and not by food intake *per se*. To address this experimentally and to disassociate increased wakefulness from the energy deficit imposed by SD, additional experiments are required involving offering more palatable food or artificial feeding during SD.

In summary, our study showed that in mice a one-time, short sleep deprivation leads to an immediate caloric deficit that resulted in an immediate loss of body weight. This finding contrasts observations in humans documenting an increase in food intake during sleep deprivation and associated weight gain. The notion that in mice a sleep deprivation also represents a food deprivation puts the consequences of this intervention, which are commonly attributed to the loss of sleep, in a new perspective. For example, recovery sleep in mice could therefore not only be seen as a response to correct for sleep time lost but also as a behavioral strategy to balance the body’s energy budget.

## Methods

### Animals and housing conditions

All mice were male C57BL/6J mice, aged 10-16 weeks at the time of recording and were individually housed during the recordings under 12h light/12h dark conditions at 23°C ambient temperature. Food (standard chow: 3436 Kliba Nafag, Switzerland with a 13.1kJ/g metabolizable energy content) and water were given *ad libitum*. All animal procedures followed the guidelines of Swiss federal law and were pre-approved by the Veterinary Office of the Canton of Vaud.

### Keeping mice awake

Sleep deprivations (SD) were performed between zeitgeber time (ZT)0; i.e., light onset, and ZT6, i.e., the mid-point of the 12h light phase. During this period, mice usually spent most of their time asleep. Mice were kept awake using the gentle handling method [55], which is somewhat of a misnomer as mice were not handled by the experimenters. Instead, mice were kept in their home cage and left undisturbed unless signs of sleep appeared. Sleep was prevented by introducing and again removing paper tissue, changing the litter, bringing a pipet in the animal’s proximity, or gentle tapping of the cage.

### Quantification of cortical gene expression

Four experimental groups of mice were used: i) control mice with *ad libitum* food and sleep (*n*=5), ii) mice kept awake (SD; ZT0-6) with free access to food (*n*=4), iii) control mice with *ad libitum* sleep but no access to food between ZT0-6 (*n*=3), and iv) mice with no access to sleep or food between ZT0-6 (*n*=4).

At ZT6, mice of all 4 groups were anesthetized with isoflurane and immediately sacrificed by decapitation. Cerebral cortex tissue was quickly dissected, immediately frozen in liquid nitrogen, and then stored at −80°C. Total mRNA were extracted from 30mg of cortical tissue using the *RNeasy Mini kit* (Qiagen) following the manufacturer’s instructions. Briefly, each sample was homogenized by pestle using 350μl of RLT buffer (supplemented with β–mercaptoethanol from Sigma-Aldrich instead of DTT) on ice and passed on a *QIAshredder* column (Qiagen). Contaminating DNA was removed with DNase I (Qiagen). RNA quantification was performed on a *Nanodrop ND1000* with a consistent 260/280 ratio between 1.8 and 2.1. Reverse transcription from 500ng of RNA was performed using the *SuperScript II Reverse Transcriptase* (Invitrogen, Carlsbad, NM, USA) following the manufacturer’s instructions. cDNA was diluted 10 times in DNAse free water. Reactions without enzyme (NEC) and without the template (NTC) were performed in parallel as negative controls.

qPCR was performed according to the Applied Biosystems protocol using a *7900HT Fast Real-time* PCR system with SDS 2.3 software (Applied Biosystems, Foster City, CA, USA). For each reaction, 7.2μl of the diluted cDNA was supplemented with 18.0μl *FastStart Universal Probe Master* 2x (Roche Diagnostic GmbH, Mannheim, Germany), 0.90μM of primer and 0.25μM of probe for a total reaction mix of 36.0μl. The qPCR was performed under standard cycling conditions (50°C for 2min, 95°C for 10min, followed by 45 cycles of 95°C for 15sec and 60°C for 1min). Each qPCR reaction was performed in triplicate. The oligonucleotide sequences used can be found in Supplementary Table S1. Primers and probes were purchased from Invitrogen or Eurogentec and all probes were dual-labelled with 5′FAM/3′BHQ as fluorophore and quencher. *Ribosomal protein S9* (*Rsp9*), *eukaryotic translation elongation factor 1A* (*Eef1a*), and *glyceraldehyde-3-phosphate dehydrogenase (Gapdh)* were found to be appropriate control genes to normalize qPCR experiments following the method proposed by Vandesomplele et al. [56]. The relative quantification of the expression of *Per1*, *Per2*, *Cry1*, *Dbp*, and *Homer1a* was calculated with a modified ΔΔCt method [57]. All stability calculations, normalization, and quantification were performed using *qbase^PLUS^ 1.5* (Biogazelle, Machelen, Belgium). For displaying purposes, gene expression levels were expressed relative to the levels reached in the two sleep control groups (i.e. groups i and iii). Gene expression in the control groups was not affected by having access to food or not (post-hoc *t*-tests: for *Per1, Per2, Cry1*, and *Dbp p*>0.47; for *Homer1a p*>0.09).

### Quantification of food intake

Food intake was recorded in two separate cohorts of mice (*n*=6 and 11, respectively). Mice of both cohorts were habituated to the recording cages for at least 11 days prior to the sleep deprivation (designated as Day 1 starting at light onset). In cohort 1, baseline food intake was measured continuously for 10 days (Days −9 to 0). On the following day (Day 1) mice were sleep-deprived between ZT0 and −6. The remaining 18h of Day 1 and Day 2 were considered recovery. On Day −2 recording had to be restarted and data of the entire day were discarded for all mice. Also, data of Day 0 had to be discarded due to technical problems. In cohort 2, the baseline was recorded for 5 days (Days −4 to 0), followed by Day 1 (6h SD + 18h recovery) and 4 more recovery days (Days 2 to 5). One mouse could only be recorded until dark-onset (ZT12) of Day 1; i.e., the SD plus the first 6h of recovery, due to technical problems. Combined the 2 cohorts spanned 10 days of baseline and 5 days of recovery (including the SD) with the following numbers of mice contributing to each day: Days −9 to 0: *n*=6, 6, 6, 6, 6, 17, 17, 11, 17, 11; Days 1-5: *n*=17/16, 16, 11, 11, 11. To construct a baseline food intake time-course, data were collapsed into hourly intervals and then averaged over the 5 days preceding SD (Days −4 to 0; i.e., for cohort 1: Days −4, −3, and −1; cohort 2: Days −4 to 0).

Food intake was quantified as food disappearance from the feeders with the *Phenomaster* feeding-monitoring system (TSE Systems GmbH, Bad Homburg, Germany), located in a dedicated room with restricted access to minimize the impact of human presence. Mice were individually kept in standard housing conditions (see above) with food and water *ad libitum* in cages equipped with rat-type feeders with large food and spillage containers to minimize maintenance during experiments. Feeder scales were calibrated according to the manufacturer’s instructions using 4 weights (range: 0-90g). The sensitivity of measurement was 10mg. Food intake was regularly monitored throughout the experiments from remote using *TeamViewer*.

Food disappearance was collected at 1min resolution as unfiltered, true feeder-weight values with the following analogue-to-digital converter (ADC) settings: *smoothing ADC*: 5s, *trial monitor observation interval*: 1s, and *Max Delta ADC*: 80. To assess the quality of raw data in-software filtering was disabled and was instead applied after acquisition and inspection of the raw data using *Excel*. Artefacts (mostly sudden positive or negative deflections related to mice touching the feeders) were removed for data points with values differing by more than −0.15 or +0.5g relative to that of the preceding data point by setting their values to that of the preceding data point, following the filter settings of the manufacturer’s software.

Food spillage not accounted for by the feeding system was assessed in a pilot experiment using special bedding (Lignocel^®^ nesting small; JRS, Germany) that, upon removal from the cage, reveals spillage at the cage bottom. Spillage was fully contained within the feeder as we found no measurable spillage on the cage bottom after the 3-day test. We estimate in-cage spillage to be <1%.

### Quantification of sleep

Previously published sleep-wake data [21, 22] were used to relate hourly mean values of time-spent-awake to hourly food intake values measured in the current study. Male C57BL/6J mice 10-12 weeks (*n*=12) at the time of SD were kept under our standard housing conditions (see above). EEG/EMG was recorded continuously from 4 days; 2 baseline days, followed by a 6h SD starting at ZT0 and 42h of recovery (Days 1 and 2). In 6 of these mice, recovery was recorded for an additional 5 days.

To determine sleep-wake states mice were equipped with chronic EEG and EMG electrodes under deep anesthesia according to methods detailed in [55]. The electrode leads were connected to a socket fixed to the skull to which the recording lead could later be fitted. The head assembly prevented easy access to food in the feeder and thus mice in which food-intake was monitored the EEG/EMG could not be recorded at the same time. A Xylazine (10mg/kg)/Ketamine (100mg/kg) mix was injected IP ensuring a deep plane of anesthesia for the duration of the surgery (ca. 30-40min). Analgesia was provided the evening prior and the 3 days after surgery (Dafalgan in drinking water; 200–300mg/kg). Mice were allowed to recover for at least 10 days prior to baseline recordings. Electrophysiological signals were captured and ADC at (sampling rate 2000Hz, down-sampled and stored at 200Hz) using *Embla A10* and *Somnologica-3* hard- and software, respectively (Medcare Flaga; Thornton). The sleep-wake states REM sleep, NREM sleep, and wakefulness were annotated for consecutive 4s time-windows based on the EEG and EMG patterns according to established criteria [55]. 4s wake scores were collapsed into hourly values and expressed as min/h. The two baselines were averaged to generate a single 24h time-course.

### Quantification of body weight and body composition

Male C57BL/6J mice, aged 13-14 weeks, were kept under our standard housing conditions (see above). The effect of SD on body weight was assessed in two cohorts (*n*=5 and 4, respectively). Each animal was individually housed 10 days before SD and weighed between ZT6.0-6.5 on 3 days; i.e., 4 days and 1 day before the SD and on the SD day itself. SD was scheduled from ZT0-6 as in all other experiments. The first 2 measurements were averaged and considered as baseline to which body weight after the SD was contrasted within individual mice. The two baseline measures did not differ (−0.14±0.10g; two-sided paired *t*-test, *p*=0.21, *n*=9). For weighing, animals were transported and measured while in a cardboard tube to reduce handling stress. Balance gave weights at 0.01g precision.

Following the same experimental protocol used for the body weight assessment explained above, in a 3^rd^ cohort (male C57BL/6J mice, aged 13-14 weeks, *n*=6), body composition was assessed using an EchoMRI™ 3-in-1 (Houston, Texas, US). Body weight, fat mass, lean body mass, and free and total water content were quantified. Hydration percentages were calculated as follows: (total – free water)*100/lean body mass. Mice were lightly anesthetized with isoflurane before putting them into the analyzer. The whole procedure (weighing, induction of anesthesia, data acquisition) did not take more than 5 min. Measures of body weight, and body composition (% hydration, fat mass, and free and total water) did not differ between the two baseline measures (*p*=0.29, 0.59, 0.65, 0.77, and 0.06, respectively), with the exception of lean body mass (−0.41±0.11g, 2^nd^ *vs*. 1^st^ baseline day; 2-sided paired *t*-tests, *p*=0.014, *n*=6).

### Statistics

For data organization, filtering, and collapsing into hourly values *Excel* and *TMT Pascal* (version 5.01) were used. Statistics were performed in *SAS* (version 9.4), linear and non-linear regression analyses in *SigmaPlot* (version 12.5). *SigmaPlot* was also used to generate the graphs. As the threshold of significance *α*=0.05 was used. Results are given as mean values ± SEM. Details on a number of biological (and technical) replicates per experiment are stated in Methods. Gene expression was analyzed first with 2-way analysis of variance (ANOVA) with factors ‘Food’ and ‘SD’. Post-hoc contrasts were analyzed 2-sided *t*-tests. Effects of SD on daily food-intake during recovery days (Days 1 to 5) compared to baseline (Days −4 to 0) was analyzed with a 2-way ANOVA with factors ‘Day’ and ‘SD’ followed by post-hoc paired 2-sided *t*-tests assessing deviations from the individual baseline means. Food-intake differences between corresponding accumulated hourly values of SD and recovery (Days 1 to 5) and baseline (mean Days −4 to 0) were analyzed with paired 2-sided *t*-tests as were the effects of SD on body weight and body composition.

## Acknowledgments

We thank members of the members of the Franken Lab for help with the sleep deprivations.

## Author contributions

Conceptualization: ND, FLS, PF; Methodology: ND, FLS, FP; Formal Analysis: ND, FLS, FP, PF; Investigation: ND, FLS, YE, GN; Writing – Original Draft Preparation: ND, PF; Writing – Review & Editing: ND, FLS, YE, GN, FP, PF; Visualization: PF; Supervision: PF; Project Administration: FP, PF; Funding Acquisition: PF.

## Institutional Review Board Statement

The study was conducted according to the guidelines of the Declaration of Helsinki and all animal procedures followed the guidelines of Swiss federal law and were pre-approved by the Veterinary Office of the Canton of Vaud (VD2348, VD3201×1a).

## Data Availability Statement

Data are available upon request.

---

**Figure S1:**
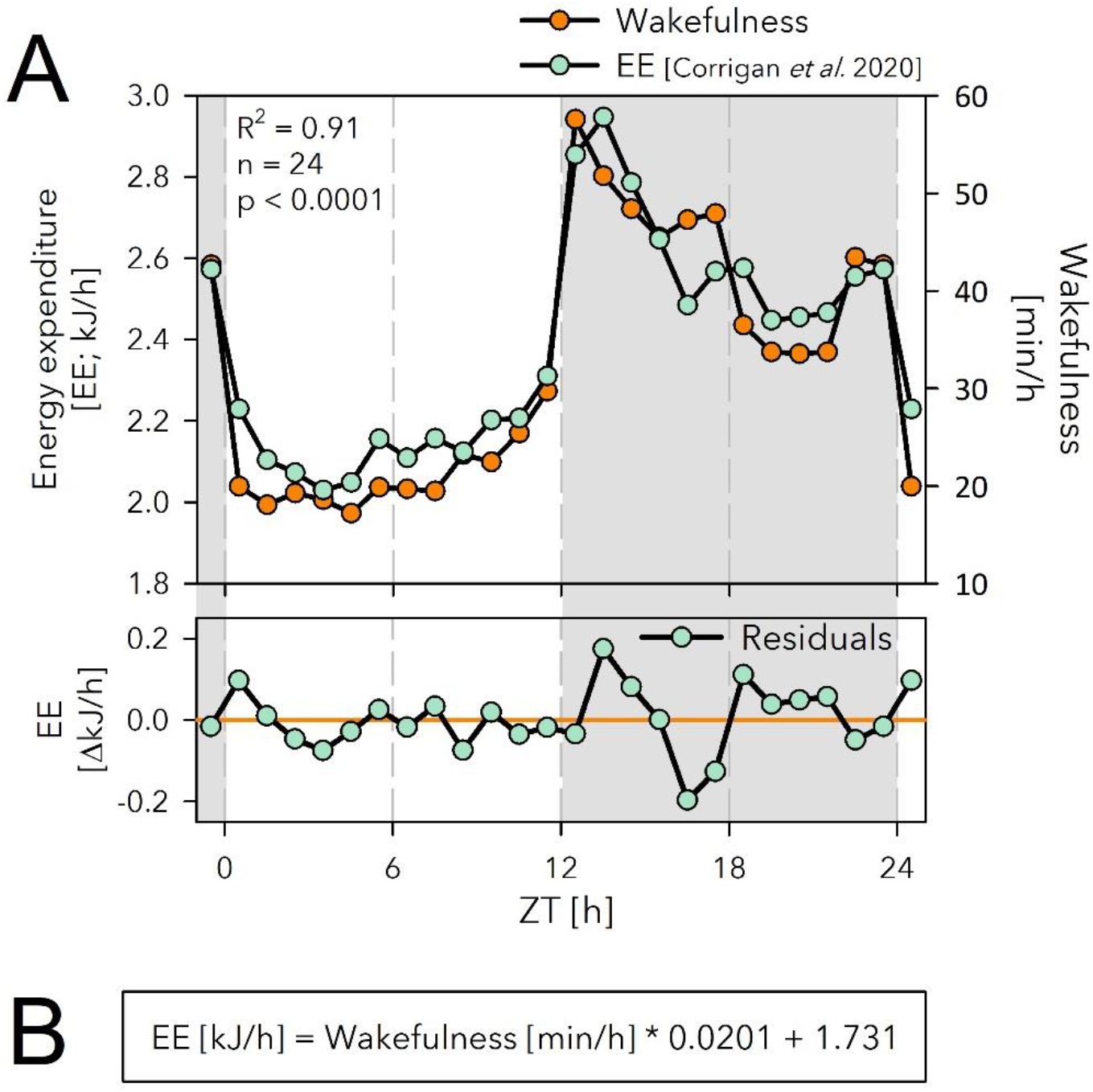
Relationship between energy expenditure (EE) and time-spent-awake under baseline conditions. (**A**) The time course of hourly values of EE (mint symbols; mean over 3 days; *n*=8, body weight: 28.4g, C57BL/6J males; data taken from Corrigan *et al*. 2020) closely follows that of time-spent-awake (orange symbols, same as in Fig. 3A). The two variables strongly correlated (linear regression) with only small residuals that did not systematically vary with time-of-day (lower graph). (**B**) Equation of the linear regression, which predicts that a mouse will spent 1.73 or 2.9kJ, if asleep or awake for 60min, respectively, in a given hour.

**Table S1:**
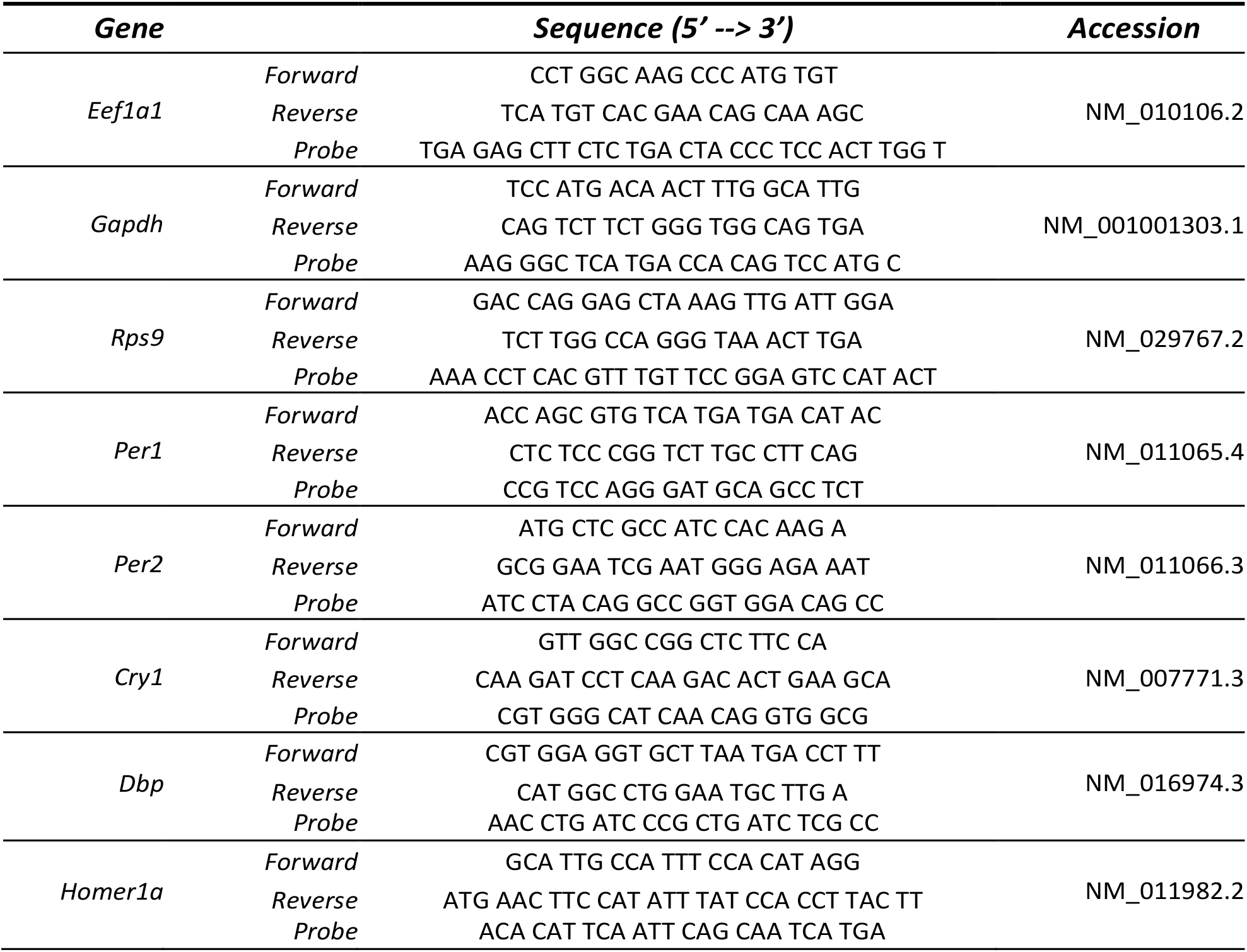
Primers and probes used for quantitative PCR.

## References

1. Krieger DT. Food and Water Restriction Shifts Corticosterone, Temperature, Activity and Brain Amine Periodicity. Endocrinology. 1974;95(5):1195–201.

2. Honma KI, Honma S, Hiroshige T. Feeding-associated corticosterone peak in rats under various feeding cycles. American Journal of Physiology-Regulatory, Integrative and Comparative Physiology. 1984;246(5):R721–R6.

3. Stephan FK. The “Other” Circadian System: Food as a Zeitgeber. Journal of Biological Rhythms. 2002;17(4):284–92.

4. Mistlberger R. Food as circadian time cue for appetitive behavior [version 1; peer review: 5 approved]. F1000Research. 2020;9(61).

5. Damiola F, Le Minh N, Preitner N, Kornmann B, Fleury-Olela F, Schibler U. Restricted feeding uncouples circadian oscillators in peripheral tissues from the central pacemaker in the suprachiasmatic nucleus. Genes Dev. 2000;14(23):2950–61.

6. Stokkan KA, Yamazaki S, Tei H, Sakaki Y, Menaker M. Entrainment of the circadian clock in the liver by feeding. Science. 2001;291(5503):490–3.

7. Wisor JP, O’Hara BF, Terao A, Selby CP, Kilduff TS, Sancar A, et al. A role for cryptochromes in sleep regulation. BMC Neurosci. 2002;3:20.

8. Curie T, Maret S, Emmenegger Y, Franken P. In Vivo Imaging of the Central and Peripheral Effects of Sleep Deprivation and Suprachiasmatic Nuclei Lesion on PERIOD-2 Protein in Mice. Sleep. 2015;38(9):1381–94.

9. Franken P. A role for clock genes in sleep homeostasis. Current Opinion in Neurobiology. 2013;23(5):864–72.

10. Franken P, Thomason R, Heller HC, O’Hara BF. A non-circadian role for clock-genes in sleep homeostasis: a strain comparison. BMC neuroscience. 2007;8:87-.

11. Hoekstra MM, Jan M, Katsioudi G, Emmenegger Y, Franken P. The sleep-wake distribution contributes to the peripheral rhythms in PERIOD-2. Elife. 2021;10.

12. Spiegel K, Tasali E, Penev P, Cauter EV. Brief communication: sleep curtailment in healthy young men is associated with decreased leptin levels, elevated ghrelin levels, and increased hunger and appetite. Annals of internal medicine. 2004;141(11):846–50.

13. Fang Z, Spaeth AM, Ma N, Zhu S, Hu S, Goel N, et al. Altered salience network connectivity predicts macronutrient intake after sleep deprivation. Scientific Reports. 2015;5(1):8215.

14. Jung CM, Melanson EL, Frydendall EJ, Perreault L, Eckel RH, Wright KP. Energy expenditure during sleep, sleep deprivation and sleep following sleep deprivation in adult humans. The Journal of Physiology. 2011;589(1):235–44.

15. Maret S, Dorsaz S, Gurcel L, Pradervand S, Petit B, Pfister C, et al. Homer1a is a core brain molecular correlate of sleep loss. Proc Natl Acad Sci U S A. 2007;104(50):20090–5.

16. Mackiewicz M, Paigen B, Naidoo N, Pack AI. Analysis of the QTL for sleep homeostasis in mice: Homer1a is a likely candidate. Physiological Genomics. 2008;33(1):91–9.

17. Curie T, Mongrain V, Dorsaz S, Mang GM, Emmenegger Y, Franken P. Homeostatic and circadian contribution to EEG and molecular state variables of sleep regulation. Sleep. 2013;36(3):311–23.

18. Scharf MT, Naidoo N, Zimmerman JE, Pack AI. The energy hypothesis of sleep revisited. Progress in Neurobiology. 2008;86(3):264–80.

19. Benington JH, Craig Heller H. Restoration of brain energy metabolism as the function of sleep. Progress in Neurobiology. 1995;45(4):347–60.

20. Vyazovskiy VV, Cirelli C, Tononi G, Tobler I. Cortical metabolic rates as measured by 2-deoxyglucose-uptake are increased after waking and decreased after sleep in mice. Brain Research Bulletin. 2008;75(5):591–7.

21. Diessler S, Jan M, Emmenegger Y, Guex N, Middleton B, Skene DJ, et al. A systems genetics resource and analysis of sleep regulation in the mouse. PLoS biology. 2018;16(8):e2005750.

22. Hor CN, Yeung J, Jan M, Emmenegger Y, Hubbard J, Xenarios I, et al. Sleep–wake-driven and circadian contributions to daily rhythms in gene expression and chromatin accessibility in the murine cortex. Proceedings of the National Academy of Sciences. 2019;116(51):25773–83.

23. Blaxter K. Energy metabolism in animals and man: CUP Archive; 1989.

24. Terpstra AHM. Differences between Humans and Mice in Efficacy of the Body Fat Lowering Effect of Conjugated Linoleic Acid: Role of Metabolic Rate. The Journal of Nutrition. 2001;131(7):2067–8.

25. Corrigan JK, Ramachandran D, He Y, Palmer CJ, Jurczak MJ, Chen R, et al. A big-data approach to understanding metabolic rate and response to obesity in laboratory mice. eLife. 2020;9:e53560.

26. Guo J, Hall KD. Predicting Changes of Body Weight, Body Fat, Energy Expenditure and Metabolic Fuel Selection in C57BL/6 Mice. PLoS One. 2011;6(1):e15961.

27. Ayala JE, Bracy DP, McGuinness OP, Wasserman DH. Considerations in the Design of Hyperinsulinemic-Euglycemic Clamps in the Conscious Mouse. Diabetes. 2006;55(2):390–7.

28. Champy MF, Selloum M, Piard L, Zeitler V, Caradec C, Chambon P, et al. Mouse functional genomics requires standardization of mouse handling and housing conditions. Mammalian Genome. 2004;15(10):768–83.

29. Green CL, Mitchell SE, Derous D, Wang Y, Chen L, Han J-DJ, et al. The effects of graded levels of calorie restriction: IX. Global metabolomic screen reveals modulation of carnitines, sphingolipids and bile acids in the liver of C57BL/6 mice. Aging Cell. 2017;16(3):529–40.

30. Mongrain V, Hernandez SA, Pradervand S, Dorsaz S, Curie T, Hagiwara G, et al. Separating the Contribution of Glucocorticoids and Wakefulness to the Molecular and Electrophysiological Correlates of Sleep Homeostasis. Sleep. 2010;33(9):1147–57.

31. Seugnet L, Boero J, Gottschalk L, Duntley SP, Shaw PJ. Identification of a biomarker for sleep drive in flies and humans. Proceedings of the National Academy of Sciences. 2006;103(52):19913–8.

32. Jensen TL, Kiersgaard MK, Sørensen DB, Mikkelsen LF. Fasting of mice: a review. Lab Anim. 2013;47(4):225–40.

33. Sato N, Marui S, Ozaki M, Nagashima K. Cold exposure and/or fasting modulate the relationship between sleep and body temperature rhythms in mice. Physiology & Behavior. 2015;149:69–75.

34. Marcheva B, Ramsey KM, Affinati A, Bass J. Clock genes and metabolic disease. Journal of Applied Physiology. 2009;107(5):1638–46.

35. Panda S, Antoch MP, Miller BH, Su AI, Schook AB, Straume M, et al. Coordinated transcription of key pathways in the mouse by the circadian clock. Cell. 2002;109(3):307–20.

36. Fifel K, Meijer JH, Deboer T. Long-term effects of sleep deprivation on neuronal activity in four hypothalamic areas. Neurobiol Dis. 2018;109(Pt A):54–63.

37. Berthoud HR, Münzberg H. The lateral hypothalamus as integrator of metabolic and environmental needs: from electrical self-stimulation to opto-genetics. Physiol Behav. 2011;104(1):29–39.

38. Bodosi B, Gardi J, Hajdu I, Szentirmai E, F. Obal J, Krueger JM. Rhythms of ghrelin, leptin, and sleep in rats: effects of the normal diurnal cycle, restricted feeding, and sleep deprivation. American Journal of Physiology-Regulatory, Integrative and Comparative Physiology. 2004;287(5):R1071–R9.

39. Dudley CA, Erbel-Sieler C, Estill SJ, Reick M, Franken P, Pitts S, et al. Altered patterns of sleep and behavioral adaptability in NPAS2-deficient mice. Science. 2003;301(5631):379–83.

40. Franken P, Malafosse A, Tafti M. Genetic Determinants of Sleep Regulation in Inbred Mice. Sleep. 1999;22(2):155–69.

41. Meerlo P, Sgoifo A, Suchecki D. Restricted and disrupted sleep: Effects on autonomic function, neuroendocrine stress systems and stress responsivity. Sleep Medicine Reviews. 2008;12(3):197–210.

42. Kohsaka A, Bass J. A sense of time: how molecular clocks organize metabolism. Trends in Endocrinology & Metabolism. 2007;18(1):4–11.

43. Zhang S, Zeitzer JM, Sakurai T, Nishino S, Mignot E. Sleep/wake fragmentation disrupts metabolism in a mouse model of narcolepsy. The Journal of Physiology. 2007;581(2):649–63.

44. Ko CH, Takahashi JS. Molecular components of the mammalian circadian clock. Human Molecular Genetics. 2006;15(suppl_2):R271–R7.

45. Mendoza J. Circadian Clocks: Setting Time By Food. Journal of Neuroendocrinology. 2007;19(2):127–37.

46. Albrecht U, Sun ZS, Eichele G, Lee CC. A Differential Response of Two Putative Mammalian Circadian Regulators, mper1and mper2, to Light. Cell. 1997;91(7):1055–64.

47. Balsalobre A, Brown SA, Marcacci L, Tronche F, Kellendonk C, Reichardt HM, et al. Resetting of Circadian Time in Peripheral Tissues by Glucocorticoid Signaling. Science. 2000;289(5488):2344–7.

48. Rutter J, Reick M, Wu LC, McKnight SL. Regulation of Clock and NPAS2 DNA Binding by the Redox State of NAD Cofactors. Science. 2001;293(5529):510–4.

49. Brown SA, Zumbrunn G, Fleury-Olela F, Preitner N, Schibler U. Rhythms of Mammalian Body Temperature Can Sustain Peripheral Circadian Clocks. Current Biology. 2002;12(18):1574–83.

50. Jiang W-G, Li S-X, Zhou S-J, Sun Y, Shi J, Lu L. Chronic unpredictable stress induces a reversible change of PER2 rhythm in the suprachiasmatic nucleus. Brain Research. 2011;1399:25–32.

51. Challet E, Poirel V-J, Malan A, Pévet P. Light exposure during daytime modulates expression of Per1 and Per2 clock genes in the suprachiasmatic nuclei of mice. Journal of Neuroscience Research. 2003;72(5):629–37.

52. Steinlechner S, Jacobmeier B, Scherbarth F, Dernbach H, Kruse F, Albrecht U. Robust Circadian Rhythmicity of Per1 and Per2 Mutant Mice in Constant Light, and Dynamics of Per1 and Per2 Gene Expression under Long and Short Photoperiods. Journal of Biological Rhythms. 2002;17(3):202–9.

53. Reyes BA, Pendergast JS, Yamazaki S. Mammalian peripheral circadian oscillators are temperature compensated. Journal of biological rhythms. 2008;23(1):95–8.

54. Gerber A, Esnault C, Aubert G, Treisman R, Pralong F, Schibler U. Blood-Borne Circadian Signal Stimulates Daily Oscillations in Actin Dynamics and SRF Activity. Cell. 2013;152(3):492–503.

55. Mang GM, Franken P. Sleep and EEG Phenotyping in Mice. Curr Protoc Mouse Biol. 2012;2(1):55–74.

56. Vandesompele J, De Preter K, Pattyn F, Poppe B, Van Roy N, De Paepe A, et al. Accurate normalization of real-time quantitative RT-PCR data by geometric averaging of multiple internal control genes. Genome Biology. 2002;3(7):research0034.1.

57. Hellemans J, Mortier G, De Paepe A, Speleman F, Vandesompele J. qBase relative quantification framework and software for management and automated analysis of real-time quantitative PCR data. Genome Biology. 2007;8(2):R19.

